# A mycelium biofactory: Novel biomaterial obtained by culturing *Ganoderma sessile* on a potential osteogenic substrate

**DOI:** 10.1101/2025.09.25.678536

**Authors:** Noelia Laura D’Elía, Pablo Postemsky, Javier Sartuqui, Damián Placente, Verónica Gonzalez-Pardo, Daniel Ercoli, Paula Verónica Messina

## Abstract

Oral diseases represent a significant public health challenge, particularly periodontal and peri-implant diseases that result in considerable bone loss and diminished quality of life. To address these issues, tissue engineering techniques such as guided bone regeneration (GBR) utilize barrier membranes to promote bone defect healing. This study introduces a novel membrane employing Ganoderma sessile mycelium as a microstructural director for osteogenic biomaterials, achieving a mycelium-alginate-nanohydroxyapatite (My-ALG-HA) composite. This innovative strategy addresses the limitations of conventional biomaterials, including inadequate management of chewing forces, limited blood supply and microbial contamination. This approach uses the unique hierarchical structure and hydrophobic properties of Ganoderma mycelium alongside the biocompatibility and hydrophilicity of the ALG-HA system to improve the structural integrity and biological functionality. The mycelium colonizes the ALG-HA substrate, forming a porous trabecular bone-like network. Water contact angle assays indicated an anisotropic interaction behavior, and tensile testing confirmed the material ductility. Incorporating mycelium-derived molecules contributes to its hydrophobicity and resistance to degradation in simulated physiological conditions. Additionally, the My-ALG-HA biomaterial demonstrates human blood hemocompatibility and osteoblast-like cell (MC3T3-E1) cytocompatibility, highlighting its potential as an advanced solution for bone tissue regeneration in regenerative medicine.

## INTRODUCTION

Oral diseases affect approximately 3.5 billion people worldwide in 2019, making them the most prevalent health issues globally^1^. Among these, periodontal and peri-implant diseases are inflammatory conditions caused by microbial activity that lead to the loss of bone-supporting teeth. Managing these diseases presents significant healthcare challenges that impact patients’ life quality^2^. In addition to periodontitis and peri-implantitis, other factors such as post-extraction atrophy, trauma, tumor removal, and congenital conditions like cleft lip and palate can result in deficiencies in orofacial tissues. To address these challenges, tissue engineering focuses on restoring lost or damaged tissues through techniques such as guided bone regeneration (GBR). The GBR technique uses barrier membranes to facilitate bone defect healing by preventing soft-tissue growth. GBR membranes can be non-absorbable or absorbable according to their degradation dynamics. Non-absorbable membranes, such as polytetrafluoroethylene (PTFE) and titanium, provide excellent mechanical properties but require surgical removal due to their inert nature. In contrast, absorbable membranes like collagen offer better biocompatibility but often face challenges related to degradation rates and strength^2^. Therefore, there is a need for new biomaterials that can overcome the limitations of current options, including managing chewing forces, addressing limited blood supply to tooth and implant surfaces, and controlling microbial contamination^3^.

In the search for innovative biomaterials, scientists are increasingly drawing inspiration from natural systems. This approach led to biomaterials characterized by ligand-like functional units derived from fungal products development^4^. The structural framework of these novel materials consists of fungal mycelium growing in conjunction with the substrate to form a three-dimensional network. This network acts as a fibrous component and binding agent, developing highly porous scaffolds^5^ that facilitate tissue regeneration.

Among these fungal-derived biomaterials, Ganoderma, a therapeutic genus of mushrooms with a rich history in traditional Chinese medicine shows significant biomedical potential due to its various biological effects, including anticancer, immunomodulatory, antioxidant, anti-inflammatory, and antimicrobial properties^6–8^. Researchers refer to Ganoderma as a “biofactory” due to its ability to synthesize various bioactive compounds^9^. For example, Ling Zhi-8 (LZ-8), a protein derived from Ganoderma lucidum acts as a potent immune modulator by enhancing cytokine expression, making it a promising candidate for osteoporosis treatment and prevention^10^. The bioactive components of Ganoderma work synergistically; their medicinal efficacy diminishes when isolated^11^. Thus, utilizing complete mycelium as a biomaterial represents an effective strategy for harnessing its full therapeutic potential.

Fungal mycelium possesses unique structural characteristics that are beneficial for biomaterial development. As a hierarchical structure, mycelium forms an interconnected network through hyphal aggregation, maximizing surface area for nutrient absorption and enabling efficient environmental exploitation. This colonization capacity allows mycelium to thrive on various substrates while adapting its form based on specific culturing conditions^12^.

Recent research has shown promising results for using whole fungal mycelia in tissue engineering applications. For instance, Narayanan et al. produced nanofibrillar scaffolds from Aspergillus sp. mycelium that exhibited biocompatibility with human keratinocytes and potential for tissue regeneration^13^. Additionally, polylactic acid and G. lucidum mycelium fibrils composite materials demonstrated biocompatibility with fibroblasts while promoting cell proliferation and collagen production. Regarding bone tissue regeneration, the fungal cell wall α-chitin promotes hydroxyapatite deposition and facilitates keratinocyte migration to wound sites, both critical processes for effective healing^14,15^. These findings underscore the potential of Ganoderma and its mycelium as valuable resources for developing advanced biomaterials for tissue regeneration and other biomedical applications.

As previously mentioned in GBR applications, biomaterials must exhibit adequate mechanical strength, appropriate degradability, and biocompatibility. Fungal mycelium not only meets these criteria but also offers natural hydrophobicity and antibacterial properties^16^. Moreover, biomaterials engineered with nano- and microscale hierarchical structures have shown significant potential in reducing bacterial adhesion and inhibiting biofilm formation. Nature-inspired surface architectures effectively prevent bacterial infections, offering a promising solution to the increasing problem of bacterial drug resistance^17^.

D’Elía et al. previously developed a bilayer membrane applied for GBR by crosslinking alginate (ALG) with varying concentrations of hydroxyapatite (HA). Their study reported positive cellular responses: osteoblasts proliferated on the mineral-rich side, while fibroblasts thrived on the fibrous side. Notably, the membranes significantly promoted osteoblast differentiation^18^, as previously demonstrated using the same synthetic HA nanoparticles alone^19,20^.

In this work, we developed a novel biomaterial by culturing *Ganoderma sessile* mycelium on an osteogenic substrate composed of alginate (ALG) and hydroxyapatite nanoparticles (HA). Our objective is to enhance the structural and biomedical features of these ALG-HA membranes by incorporating the fibrous framework of mycelium. By developing a mycelium-alginate-nanohydroxyapatite (My-ALG-HA) biomaterial, we aim to introduce unique structural and biological properties particularly suited for applications in bone tissue regeneration. We studied the effects on the hyphal structure of mycelium inactivation and sterilization. The My-ALG-HA biomaterial was chemically characterized using X-ray diffraction (XRD) and Fourier-transform infrared spectroscopy (FTIR). Structural properties and surface characteristics were analyzed through electronic microscopy and water contact angle measurements. Also, the biomaterial’s tensile test and stability under physiological conditions were evaluated. Finally, cytocompatibility was tested with osteoblast-like cells, and hemocompatibility was assessed using human blood samples.

## 1. METHODOLOGY

### 1.1 Synthesis of My-ALG-HA biomaterials

Despite the extensive research conducted on *Ganoderma lucidum*, the mycelium of *Ganoderma sessile* was chosen for the production of this biomaterial due to its genetic and phylogenetic relationship with *G. lucidum* and its genetic and phenotypic similarity^21,22^. Additionally, *G. sessile* is a well-known species that has been extensively studied by this research team^23^. The mycelium of *G. sessile* was inoculated on substrates formed by ALG-HA hydrogels with *in vitro* osteogenic potential^18^, to obtain biomaterials with superior biological and mechanical characteristics.

The substrate hydrogel was synthesized using a methodology developed previously^18^, with modifications. Briefly, 2 % w/v alginic acid was prepared in ddH_2_O and kept under 350 rpm stirring at 60 °C for 30 min. As a source of nutrients necessary for the mycelium’s growth, malt extract, yeast extract and sucrose were incorporated by sequence, and stirred in the same conditions until their complete solubilization, to obtain final concentrations of 2 % w/v, 0.2 % w/v and 1 % w/v respectively. A volume of 8 ml of this substrate solution was added to a 60 mm diameter Petri dish and dried at 50 °C for 24 h. After drying, the membrane formed was carefully detached and immersed in 15 ml of crosslinking solution composed of 110 mM CaCl_2_ and customized hydroxyapatite nanoparticles (HA)^24^ solutions in ddH_2_O in different concentrations: 0, 1.1, 4.4 and 8.8 % w/v. The substrates were stirred at 150 rpm, at room temperature (RT) for 4 h, to guarantee a complete ALG-HA crosslinking and then, they were sterilized by autoclave at 120 °C for 20 min. Under a laminar flow hood, a 4 mm^2^ portion of *G. sessile* mycelium, strain E47 (CERZOS-UNS-CONICET, Bahía Blanca, Argentina), was inoculated on the substrate, and incubated at 25 °C in the darkness for 12 days. Then, mycelium-substrates were frozen at -30 °C for 72 h and dehydrated for 24 h in a freeze-dryer (Rificor L-A-B4), under a pressure of 0.038 mmHg for 24 h. Finally, they were autoclaved at 120 °C for 20 min, to inactivate and sterilize the mycelium. Final biomaterials were named My-ALG, My-ALG-1.1HA, My-ALG-4.4HA, and My-ALG-8.8 HA. Biomaterials without HA served as control of the HA effect. The impact of varying HA concentrations in the mycelium colonization of the substrate over time was assessed through photographic documentation. The final My-ALG-HA biomaterial macrostructure aspect was studied by measuring the coverage area using Image J software before and after freeze-drying and autoclaving.

To corroborate the complete mycelium inactivation in the final biomaterials, a 4 mm^2^ portion of autoclaved *G. sessile* E47 mycelium was re-cultivated in MYSA culture medium (2 % w/v malt extract, 0.2 % w/v yeast extract and 1 % w/v sucrose, 2 % agar, pH = 6), and incubated at 25 °C in the darkness for 60 days. The absence of mycelium colonization of the substrate was considered as a non-viable mycelium.

### 1.2 Physicochemical analysis

X-ray Powder Diffraction (XRD) measurements were conducted using a Philips PW 1710 diffractometer equipped with a Cu Kα radiation source (λ = 1.5418 nm) and a graphite monochromator, operating at 45 kV, 30 mA, and 25 °C.

Also, previously dried and grounded biomaterials were homogenized with 1 % w/w KBr (FTIR grade, 99 %, Sigma-Aldrich, ref. n. 221864). The mixture was further dried at 60 °C for 48 h and placed in a micro-sample cup. The characterization was performed using a Fourier transform infrared spectroscopy (FTIR, NICOLET Nexus 470) with an AVATAR Smart Diffuse Reflectance accessory. The measurements were done at 2 cm^-1^ resolution and 64 scans min^-1^. The KBr spectrum was collected and subtracted from the samples’ spectra.

### 1.3 Structural analysis and surface characterization

#### 1.3.1 Transmission electron microscopy (TEM)

The cellular ultrastructure was studied to evaluate the effects of autoclaving (120 °C, 20 min) on the hyphae structure. A sample of biomaterial was immersed in a 0.067 M phosphate-buffered saline (PBS, pH = 7.2) solution, with glutaraldehyde at 2.5 %. Subsequently, it was washed three times with PBS. For fixation, 2 % OsO_4_ was added and immersed for 1 h, then washed four times with ddH_2_O for 15 min. Dehydration was carried out using a gradient of ethanol/distilled water: 30, 50 % v/v for 15 min and 70 % v/v overnight. Ethanol reached 96 %, and dehydration continued with 50 % v/v acetone/ethanol for 15 min until reaching 100 % acetone for 1 h. Then the samples were infiltrated with Spurr resin/acetone at RT: 1:3, 1:1 v/v (1 h each) and 3:1, v/v overnight. The samples continued in pure resin for a day. The polymerization of the resin was carried out at 70 °C overnight. Ultrathin sections were obtained with an ultramicrotome (LKB, 2088), deposited on 300 mesh copper grids and contrasted with uranyl acetate (1 min) and lead citrate (5 min). Transition electron microscopy (TEM) measurements were carried out on a JEOL 100CXII microscope with a nominal resolution of 0.6 nm, operated with an acceleration voltage of 80 kV, and magnification ranges of ×80000–×100000.

#### 1.3.2 Field emission scanning electron microscopy (FE-SEM)

The biomaterial was cut and mounted on stubs with double-sided tape. Gold or carbon was then evaporated using an Argon plasma metal evaporator (91000 Model 3, Pelco). Biomaterial was dispersed on an aluminium conductive adhesive tape (3M^®^) and metallized with gold or carbon in a sputter coater. The observation was performed in a variable pressure Scanning Electron Microscope (SEM) (LEO EVO 40XVP-M) equipped with SE, SEVP and BSE detectors and connected to a microanalysis system, operated at 7.0 kV with a resolution (WD) of 2.1 nm. Images of representative areas were acquired at different magnifications (25000x and 30000x). The associated energy-dispersive X-ray spectroscopy (EDS) (Oxford X-max 50) provided qualitative information about surface elemental composition.

#### 1.3.3 Water contact angle assay

The water contact angle between the biomaterial surface and a water drop was measured at RT using a tailored system described by Belén and col.^25^. Briefly, the system consists of placing biomaterial disks in sample holders aligned with a digital microscope. Subsequently, a tube was placed above the sample holder to form the drop. The other end of the tube connects to a syringe attached to a syringe pump, Thermometric 612 syringe pump 2, forming reproducible drops in all measurements. A water droplet of 2.0 μL was deposited over the disk to acquire the image taken within 3 s of liquid deposition by using the digital microscope (Andonstar digital microscope). All images were processed using ImageJ software^26^ and the contact angle measurements were carried out with an accuracy of ±1 using the Drop Snake and LB-ADSA available module^27^. The reported contact angle values are based on six to sixteen repeats.

### 1.4 Mechanical and stability properties

#### 1.4.1 Tensile Properties

The biomaterial was sliced into strips 5 cm long and 1 cm wide. Using an Instron Universal Testing Machine (Model 3369) equipped with a 1 kN load cell, the Young’s modulus (E), maximum tensile stress (σ_max_), maximum tensile strain (ε_max_), yield stress (σ_y_), and yield strain (ε_y_) were measured longitudinally. These tests were conducted at a speed of testing of 1 mm/min and an initial separation gap of 0.5 cm. The maximum tensile stress (σ_max_) was calculated by dividing the maximum load needed to fracture the biomaterial by its cross-sectional area. Each experiment was performed on six specimens.

#### 1.4.2 Gravimetric assay

A portion of 28 cm^2^ of biomaterial was placed in a pre-weighed petri dish and 5 ml of sterile 1X PBS was added to each dish to cover the biomaterials samples. The dishes were then sealed with cling film and incubated in a thermostatic bath at 37 °C for 24 h. After this period, the PBS was removed, and the biomaterials were oven-dried at 50 °C for 24 h. Each dried sample was then weighed. Data was accurately recorded, and the samples were once again submerged in 5 ml of sterile PBS. This process was repeated to achieve the following submerging times: 1, 2, 4, 7 and 14 days, and it was conducted in quadruplicate. The degradation profile was represented as a plot of weight loss (Equation 1) over time ^28^:

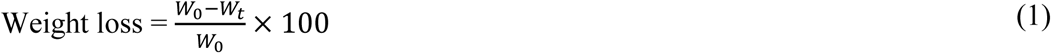

W_0_ and W_t_, are initial weight and weight at the specific time point, respectively.

### 1.5 Biocompatibility *in vitro*

#### 1.5.1 Blood sampling and percentage of hemolysis calculation

Fresh whole-blood samples were obtained from the subjects according to guidelines of Good Clinical Practices, ICH GCP Guidelines, of The Code of Ethics of the World Medical Association (the tenets of the Declaration of Helsinki) ^29–31^ Argentine National disposition ANMAT 6677/10 and Buenos Aires provincial law, no. 11044. All blood donors gave their informed consent. Neither personal information nor case details were obtained from whole-blood samples. Blood was the source of red blood cells, leukocytes, and coagulation factors. After applying a tourniquet to the patient’s upper arm, the extraction zone was disinfected, and the blood sample was obtained by venipuncture. This blood was collected in tubes and an anticoagulant was added^31^.

The hemolysis ratio (%) was assessed according to Peng and col. protocol^32^ with modifications. Briefly, 3.4 mg of biomaterials were put in contact with 700 μl of fresh entire blood and incubated at 37 °C for 24 h into Eppendorf tubes. A 3 % hydrogen peroxide solution (H_2_O_2_) was used as the positive control, and no material was used as the negative control. Supernatants of all samples were obtained by centrifugation at 3500 rpm for 5 min at RT to separate specific cell types into three layers: the top layer of plasma, the middle white layer composed of white blood cells and platelets, and the bottom red layer composed of red blood cells. Then, the supernatants were transferred into a 1 cm path-length rectangular quartz cell. The absorbance of the solutions at 414 nm was read using a UV–vis–NIR scanning spectrophotometer (Varian Cary 100 Bio). The hemolysis ratio (Equation 2) was calculated as follows:

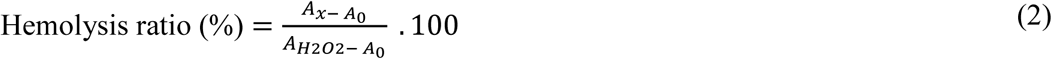

A_x_ and A_0_ were the absorbance of samples treated with different biomaterials and not treated respectively, while A_H2O2_ was the absorbance of samples treated with hydrogen peroxide solution.

#### 1.5.2 Osteoblast cells viability and morphology

As a first approach, osteoblast cell viability was assessed by crystal violet assay and cell morphology was examined by optical microscopy. MC3T3-E1 cells were cultured in flasks in Dulbecco′s Modified Eagle′s Medium (DMEM; Sigma-Aldrich St. Louis, MO, USA) supplemented with 10 % fetal bovine serum (FBS) and 1 % penicillin/streptomycin (P-S) in an incubator with a humidified atmosphere of 5 % CO_2_ in air at 37 °C. The medium was changed every other day. Upon 80 % confluence, cells were detached with a minimum amount of trypsin, which was inactivated with complete medium after 5 min. The cells were then re-cultured or used for the experiments. The biomaterial samples were washed two times with ddH_2_O for 15 min and dried at RT under sterile conditions. Then, they were mortared for 10 min in order to obtain powders and sterilized by autoclaving at 120 °C for 20 min. Afterwards, each biomaterial powder was immersed in DMEM 10 % FBS; 1 % P-S, in order to obtain a final biomaterial concentration of 1 mg/ml, and incubated at 37 °C in a water bath for 24 h while shaking at 30 rpm. MC3T3-E1 cells at passage 6 were seeded into 96-well plates at a density of 7800 cells/cm^2^ and cultured to let attach for 24 h. The next day, cell culture media was aspirated and cells were washed with 1X PBS. Then, attached cells were treated with 250 µl of 1 mg/ml biomaterial powder dispersions in complete medium, while medium without biomaterial was used as control. After 48 h of cell culture, each well was washed with PBS. For the crystal violet stain, a protocol described by Principe and col.^33^ was followed. Briefly, the cells attached to the bottom of the plate were fixed with 4 % paraformaldehyde for 7 min, stained with 0.1 % crystal violet solution for 10 min at RT and quantified by adding 0.2 % Triton x-100. Finally, the absorbance at 590 nm was measured using a microplate reader (Biotec Synergy-HT). For cell viability evaluation, the average absorbance of the control wells, which received no treatment, was set to 100 %, and the percentage of cell growth in each well was calculated relative to the control (% of control).

### 1.6 Statistical analysis

All experiments were performed in triplicate or sextuplicate, and the results are expressed as mean ± standard deviation. Statistical analysis was performed using Excel 2016 (Microsoft Corporation; USA). The t-test was used to calculate the p-value and p < 0.05 was considered significant.

## 2 RESULTS AND DISCUSSION

### 2.1 Mycelium growth and final My-ALG-HA biomaterial obtaining

To develop final My-ALG-HA biomaterials, *G. sessile* mycelium was inoculated and cultured on various substrates composed of ALG cross-linked by CaCl_2_, and different concentrations of HA: 0, 1.1, 4.4 and 8.8 % w/v. Photographic documentation revealed that the temporal progression of mycelium growth was comparable across the different substrates, with full substrate colonization observed by day 12 under all tested conditions (Figure 1). These findings suggest that the varying HA concentrations incorporated into the substrates do not significantly impact mycelium growth and colonization.

**Figure 1.**
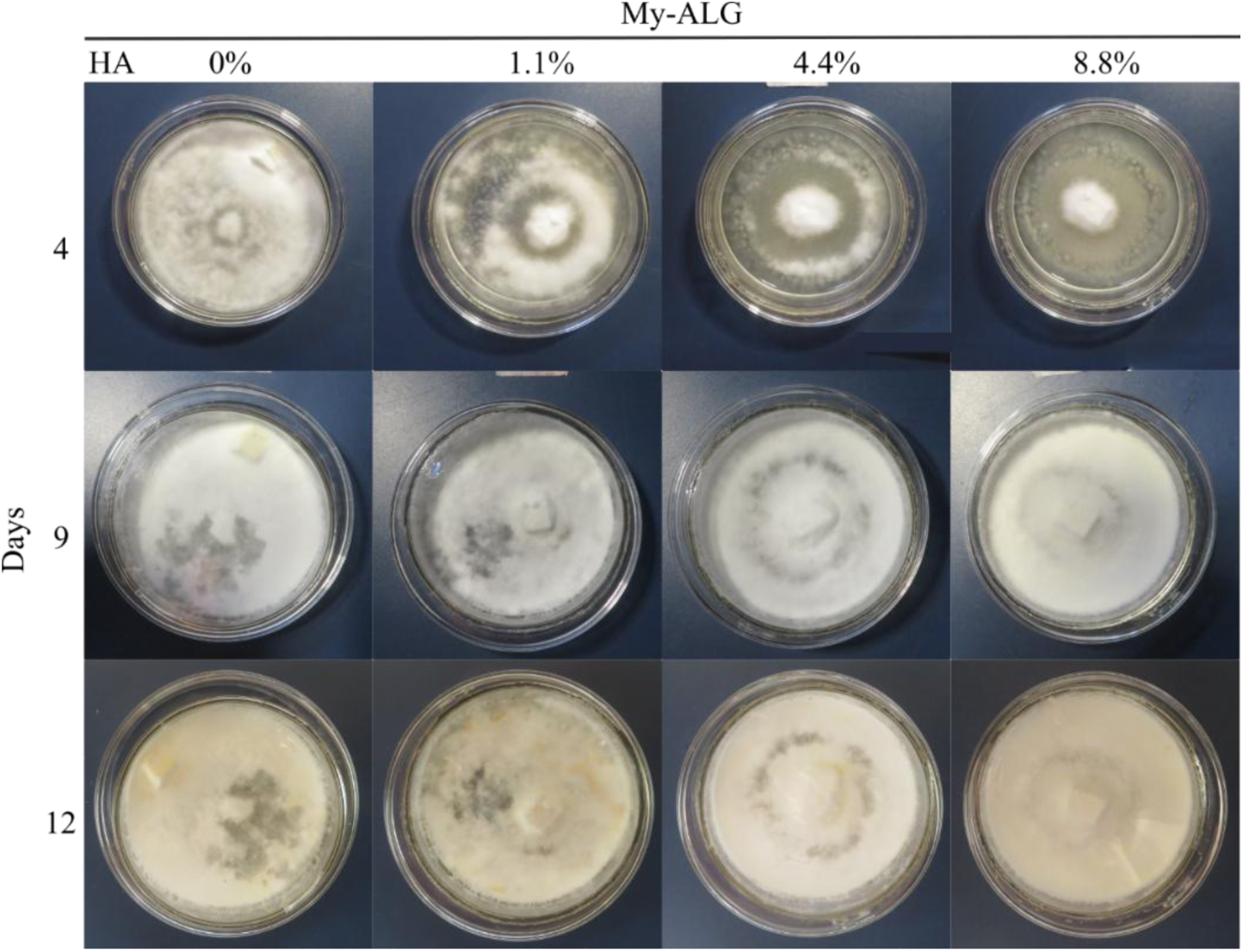
Mycelium growth on different substrates. Photographic documentation of the mycelium growth after 4, 9 and 12 culturing days in substrates composed of 2 % sodium alginate (ALG) cross-linked with CaCl_2_ only (0 % HA), and using HA in increasing concentrations 1.1, 4.4 and 8.8 % w/v.

A biomaterial intended for biomedical applications must be both sterile and inactive. Consequently, it was necessary to inactivate the mycelium cells in the final My-ALG-HA biomaterial, which was achieved by autoclaving at 120 °C for 20 min. After completing the procedure for obtaining the final My-ALG-1.1HA biomaterial, it was observed that the macrostructural characteristics were preserved, with only a 21.70% reduction in the coverage area (Figure 2).

**Figure 2.**
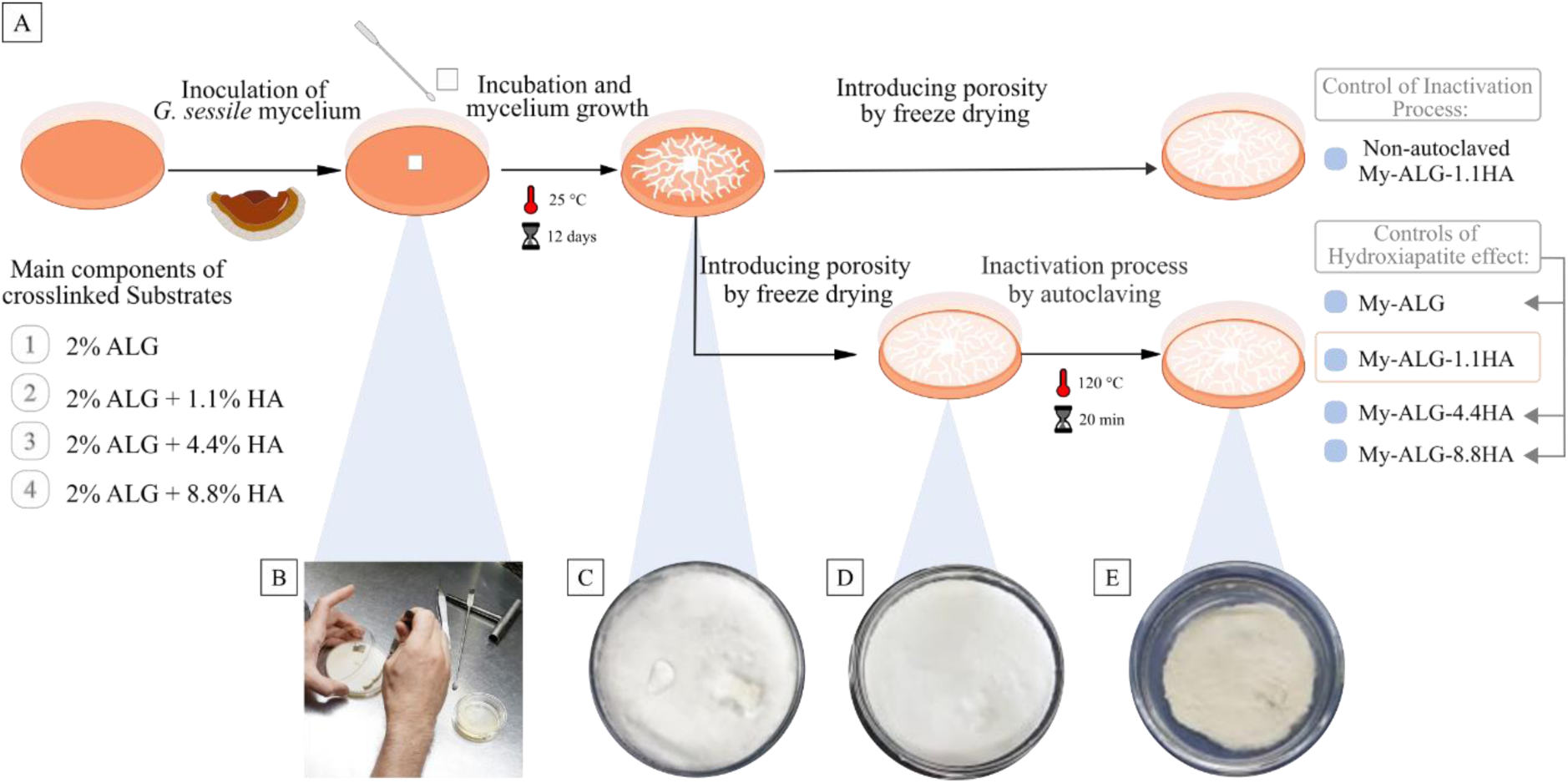
A) Procedure for obtaining My-ALG-HA biomaterials and, photographic documentation of the procedure on different stages: B) Mycelium inoculation on substrate composed of 2% alginate (ALG) and 1.1% hydroxyapatite (HA), C) mycelium cultured for 12 days at 25 °C, D) biomaterial My-ALG-1.1HA after introducing porosity by freeze-drying and, E) My-ALG-1.1HA biomaterial after inactivation process by autoclaving at 120 °C for 20 min.

The inactivation was assessed by re-incubating a fragment of the treated biomaterial on MYSA culture media. The re-growth assay showed that, following these treatments, the mycelium exhibited no detectable biological activity after a prudential time of culturing (Figure SM 1).

### 2.5.3.2 Physicochemical analysis

To characterize the crystalline phases present in the My-ALG-HA biomaterial, X-ray diffraction (XRD) analysis was performed and compared with pure synthetic hydroxyapatite (HA) nanoparticles. As expected, the XRD spectrum of My-ALG-HA displayed diffraction peaks (002) and (211), consistent with those of synthetic HA (Figures 3 A and B). Hydroxyapatite is a substrate key component and, therefore, of the final biomaterial. Additionally, the spectrum showed diffraction peaks at 24.18° and 37.35°, associated with the crystalline planes (0022) and (1032) of calcium oxalate dihydrate crystals, respectively ^34^ (Figure 3 A). The formation of calcium oxalate by fungi serves as a calcium reservoir, modulating phosphate availability in their environment. Specifically, *Ganoderma* mycelium produces calcium oxalate crystals when cultured in a Ca²⁺-enriched medium. In this regard, weddellite is the predominant form, characterized by a dehydrated tetragonal atomic arrangement^35^. Interestingly, calcium oxalate crystals serve as common binding sites for bone proteins, such as osteopontin^36^. This protein plays several critical roles in bone physiology, including promoting osteoblast adhesion and migration^37^. Consequently, the presence of calcium oxalates in this biomaterial may facilitate the adhesion of bone proteins and act as nucleation centers for the formation of additional hydroxyapatite nanocrystals *in situ*. Thus, this potential interaction warrants further investigation in future studies.

**Figure 3.**
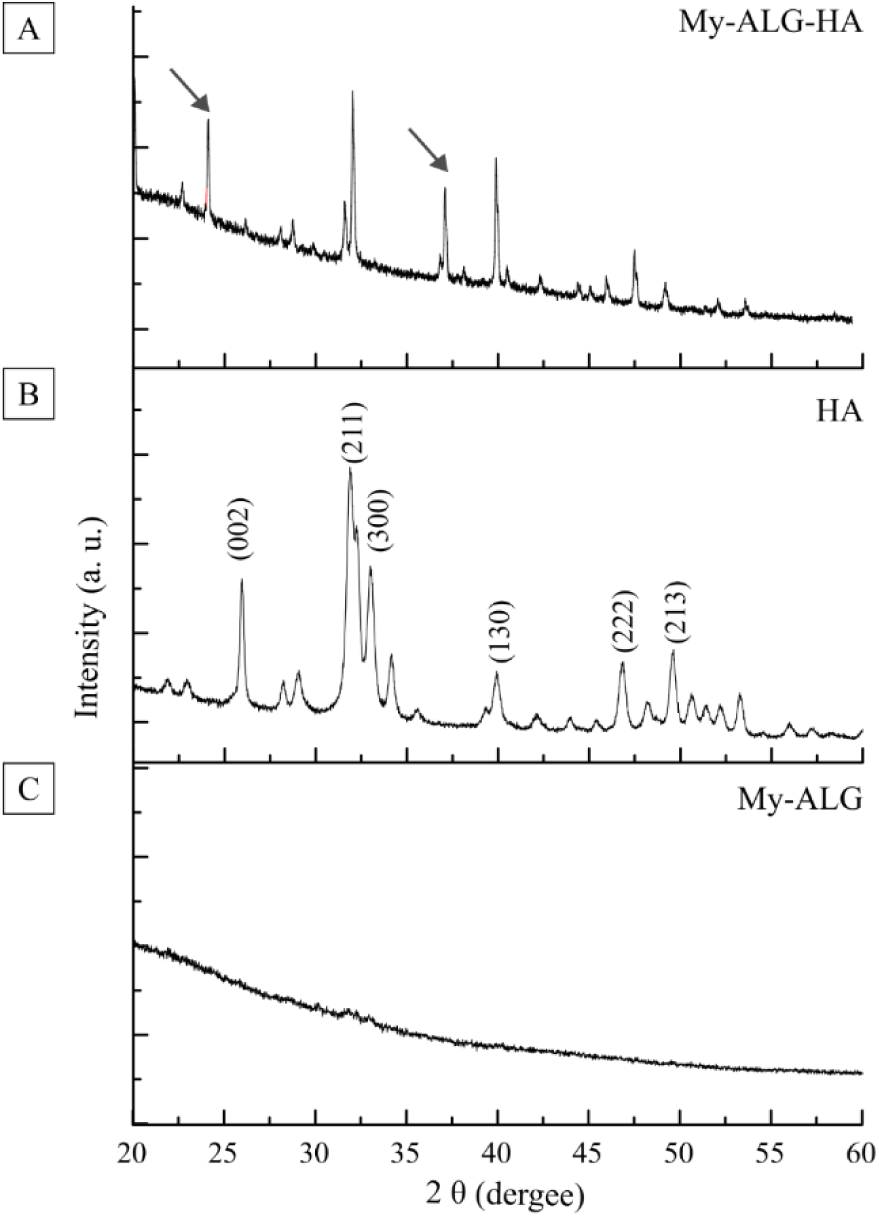
Physicochemical analysis by X-ray diffraction (XRD) spectrum. A) My-ALG-HA biomaterial. B) Pure crystalline hydroxyapatite nanoparticles (HA). C) Biomaterial without HA (My-ALG). Arrows indicate peaks corresponding to calcium oxalate dihydrate crystals.

The amorphous background observed in the My-ALG-HA spectrum can be attributed to the amorphous components of the mycelium, which were also evident in the My-ALG spectrum (Figure 3 C). No significant differences were observed among the XRD spectra of biomaterials with varying HA amounts (My-ALG-1.1HA, My-ALG-4.4HA and My-ALG-8.8HA) (Figure SM 2).

The Fourier transform infrared spectroscopy (FTIR) technique was used to analyze the composition and chemical interaction between the different components of the biomaterial. The FTIR spectrum of the My-ALG-HA biomaterial showed the presence of diverse functional groups from mycelium, ALG and HA molecules (Figure 4). Due to the diversity of functional groups present, the characteristic bands are listed in Table 1. No differences were found among FTIR spectra of My-ALG-1.1HA, My-ALG-4.4HA and My-ALG-8.8HA (Figure SM 3).

**Figure 4.**
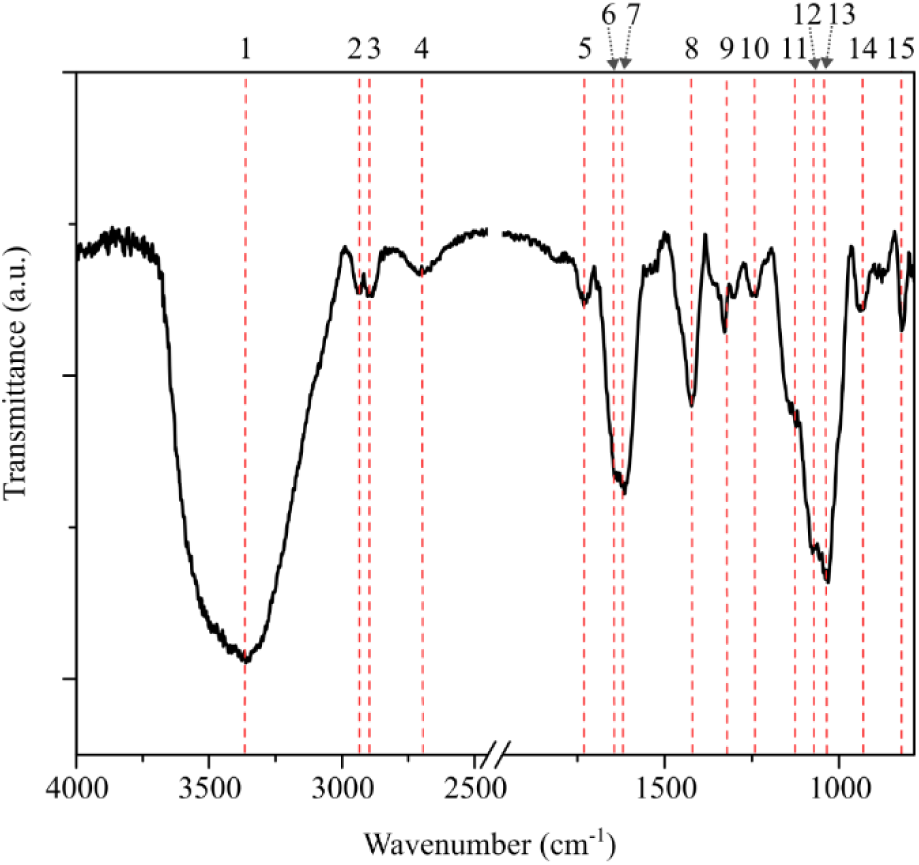
Physicochemical characterization using FTIR spectra of My-ALG-HA biomaterial. Bands indicated with numbers can be assigned to functional groups listed in Table 1.

**Table 1.** Main absorption bands (cm^-1^) in the FTIR spectrum of the My-ALG-HA biomaterial.

|  | Wavenumber (cm<sup>-1</sup>) | Assignment | My-ALG-HA component | Ref. |
| --- | --- | --- | --- | --- |
| 1 | 3365 | O–H stretching | Polysaccharides, alginate | 14,34 |
| 2 | 3055 | =C–H asymmetric stretching | Lipids | 35 |
| 3 | 2930 - 2900 | C–H stretching of alkyls | Lipids, polysaccharides | 34,36 |
| 4 | 2700 | O–H dimer stretching | Polysaccharides, alginate | 37 |
| 5 | 1730 | Ester C = O stretching | Lipids, ganoderic acids | 34 |
| 6 | 1666 | Amide I stretching (C = O stretching) | Proteins, alginate | 34,36 |
| 7 | 1620 | Amide II stretching (C–N stretching) | Proteins | 14 |
| 8 | 1421 | O–H stretching/bending in C–OH | Polysaccharides, alginate | 38 |
| 9 | 1320 | O–H bending | Polysaccharides | 38 |
| 10 | 1240 | Amide III stretching<br>(C–N, N–H stretching) | Proteins | 14,38 |
| 11 | 1153 | C–O–C asymmetric stretching | Polysaccharides | 36 |
| 12 | 1069 | C–O stretching | Polysaccharides, alginate | 14,34 |
| 13 | 1043 | C–C stretching | Polysaccharides, alginate | 14,34 |
| 14 | 927 | α-1,4-D-glucan | Polysaccharides | 39 |
| 15 | 818 | Mannan band | Polysaccharides | 34 |

Research suggests that polysaccharides and triterpenes are the most active compounds found in *Ganoderma*. These compounds can synergistically enhance many biological activities ^11^. The main bands observed in the FTIR spectrum at 1666 cm^-1^ and 1620 cm^-1^ are characteristic of functional groups of proteins, while bands at 1421 cm^-1^, 1069 cm^-1^ and 1043 cm^-1^, are typical of polysaccharides (Figure 4 and Table 1). These findings indicate that many of these molecules persisted after inactivation and sterilization treatments.

### 3.3 Structural analysis and surface characterization

The observation of hyphae ultrastructure via TEM is in accordance with the results of the inactivation assay. When examining a transversal section of the hyphae present in the non-autoclaved biomaterial (Figure 5A) compared to the autoclaved material (Figure 5B), a loss of structural integrity is observed in the latter. However, the thickness of the hyphal cell wall showed no significant differences before and after autoclaving, measuring 76.63 nm ± 15.19 nm and 77.14 nm ± 34.67 nm, respectively (Figure SM 4A). These results suggest that the cell walls were preserved in the final sterile and inactivated My-ALG-HA biomaterial. Additionally, HA nanoparticles may be observed as part of the biomaterial, situated externally to the hyphae (Figure 5C). Their presence is corroborated by the previously conducted X-ray diffraction (XRD) analysis, as well as by TEM observations, which confirm that their morphology aligns with the isolated HA used in the synthesis of the biomaterial (Figures SM 4 C and D).

**Figure 5.**
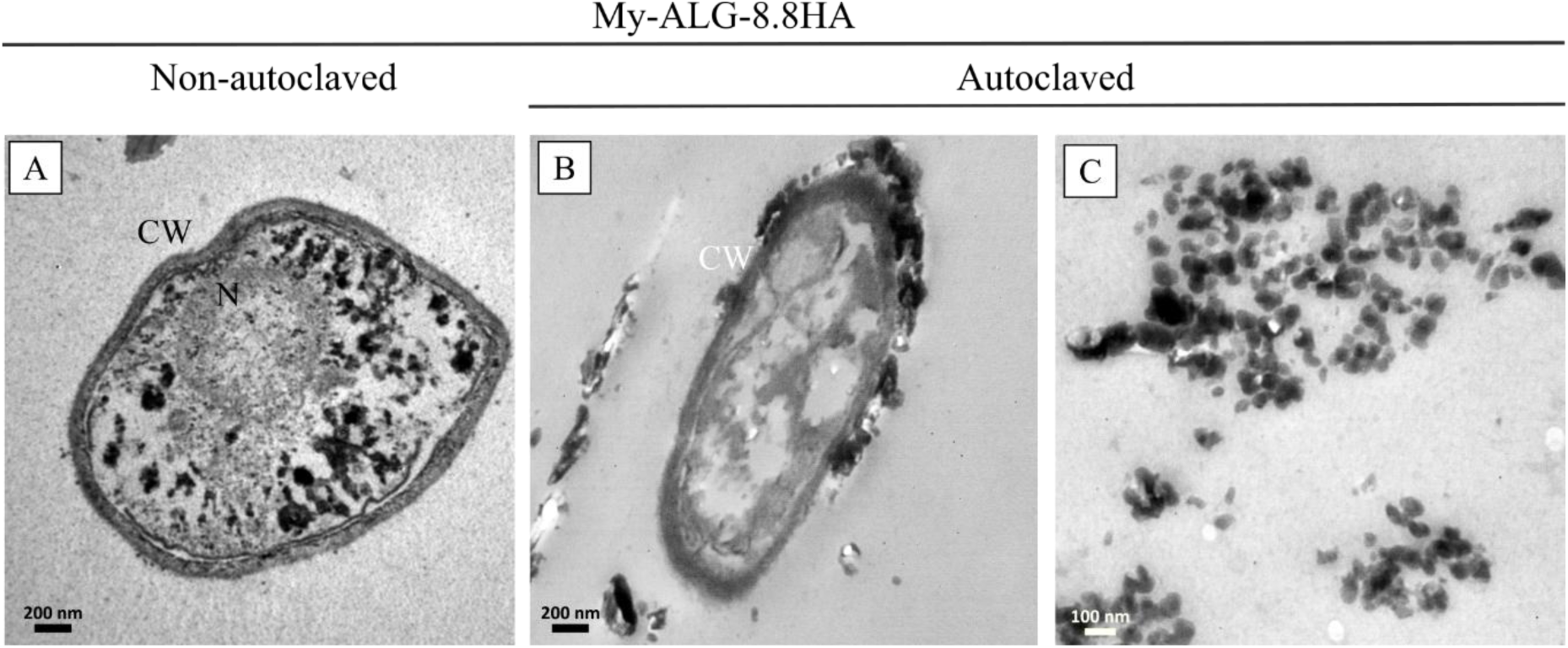
TEM micrographs of My-ALG-8.8HA biomaterial. A) Hyphae ultrastructure before and B) after freeze-drying and autoclaving. C) Section of the final My-ALG-HA biomaterial where HA are observed outside the hyphae. N: nucleus. CW: cell wall.

Customizing the surface properties of the biomaterial can enhance its biological response, including cell attachment and migration. Scanning electron microscopy (SEM) micrographs showed the mycelium acting as a biomaterial microstructure director. The mycelium colonized the ALG-HA substrate and led to the formation of a porous trabecular bone-like network with a hierarchical structure (Figure 6 and Figure SM 5). The resultant network structure is interesting as interconnected pores are essential for vital cell functions such as nutrient and oxygen supply and waste removal ^38^.Moreover, while no mineral structures were observed in the My-ALG biomaterial (Figure 6A), two distinct types of mineral structures were identified in My-ALG-1.1HA and My-ALG-8.8HA (Figures 6 B-G). The first type consists of irregular-shaped structures, which could be attributed to HA (Figure 6 D). The second type of mineral structure observed consists of tetragonal-shaped formations, likely corresponding to the “envelope-shaped” morphology typical of calcium oxalate dihydrate crystals^39^ (Figure 6 E). The presence of both hydroxyapatite and calcium oxalate dihydrate crystals was further confirmed by XRD.

**Figure 6.**
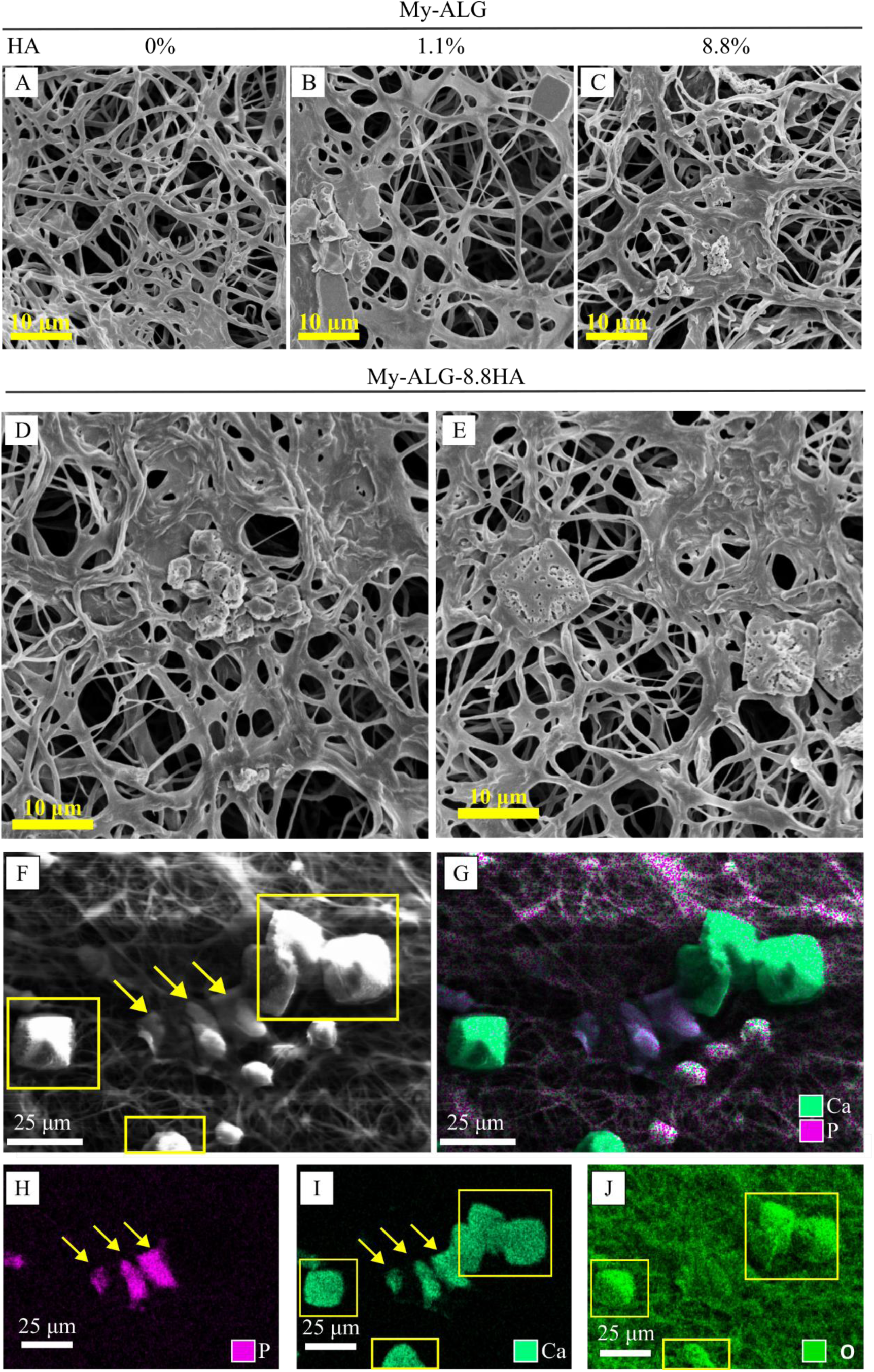
SEM micrographs of the biomaterials: A) My-ALG, B) My-ALG-1.1HA and, C) My-ALG-8.8HA. D-E) Magnification of the mineral structures on My-ALG-8.8HA biomaterial. F) Backscattered electron scanning microscopy (BSE-SEM) of the mineral structures on My-ALG-8.8HA. G) Co-localization of calcium and phosphorous atoms, H) localization of phosphorus, I) calcium, and J) oxygen atoms analyzed by energy dispersion x-ray (EDS) microanalysis. Arrows indicate the irregular-shaped and squares the tetragonal-shaped mineral structures.

To determine the elemental composition of the mineral structures observed, scanning electron microscopy-energy dispersive X-ray analysis (SEM-EDS) was employed. The co-localization of calcium and phosphate atoms in the irregular-shaped minerals could be attributed to hydroxyapatite (HA) nanoparticles, as indicated by the arrows in figures 6G to 6I (Figure 6G-I). In contrast, the tetragonal-shaped mineral structures exhibit a notable absence of phosphorus (Figure 6H), while calcium is present, along with a higher concentration of oxygen atoms, as indicated by the squares in figures 6I and 6J (Figure 6I and J). Calcium and oxygen elements are characteristic of calcium oxalate crystals.

Surface wettability, defined by hydrophobicity or hydrophilicity, is crucial for interactions between the biomaterial and its biological environment. This property influences ion interactions, protein adsorption and cell adhesion. The wettability of a surface is often expressed in terms of contact angle (CA), the angle at which a liquid-vapor interface meets the solid surface in an air environment. A water CA greater than 65° indicates hydrophobic surfaces, while angles less than 65° suggest hydrophilic surfaces ^40^. Wettability assays showed that My-ALG-HA biomaterial has different water CA depending on the side tested (Figure 7). The side where the mycelium was initially inoculated, named “mycelium-rich side”, is hydrophobic showing water CA of 143.10° ± 5.18° (Figure 7 A) and 147.77° ± 9.10° (Figure 7 A, C and E) before and after freeze-drying and autoclaving, respectively. This low wettability is expected due to the presence of hydrophobic mannoproteins and hydrophobins, in the outermost layer of the fungal cell wall. In other research, Antinori and col. found that *Ganoderma* mycelium exhibits a highly hydrophobic nature with water contact angles greater than 120° ^14^. In this sense, the micrometric roughness of the fibrous mycelium-based biomaterial strongly contributes to the hydrophobic character^41^. However, at the “alginate-rich side”, the biomaterial exhibits water CA of 36.54° ± 5.38° and 38.37° ± 1.72°, before and after freeze-dry and autoclaving respectively (Figure 7 B, D and E). This hydrophilicity could be due to the higher amount of alginate molecules, compared to mycelium, exposed on the surface. Water contact angle assays demonstrated the anisotropic water-interaction behavior of the biomaterial.

**Figure 7.**
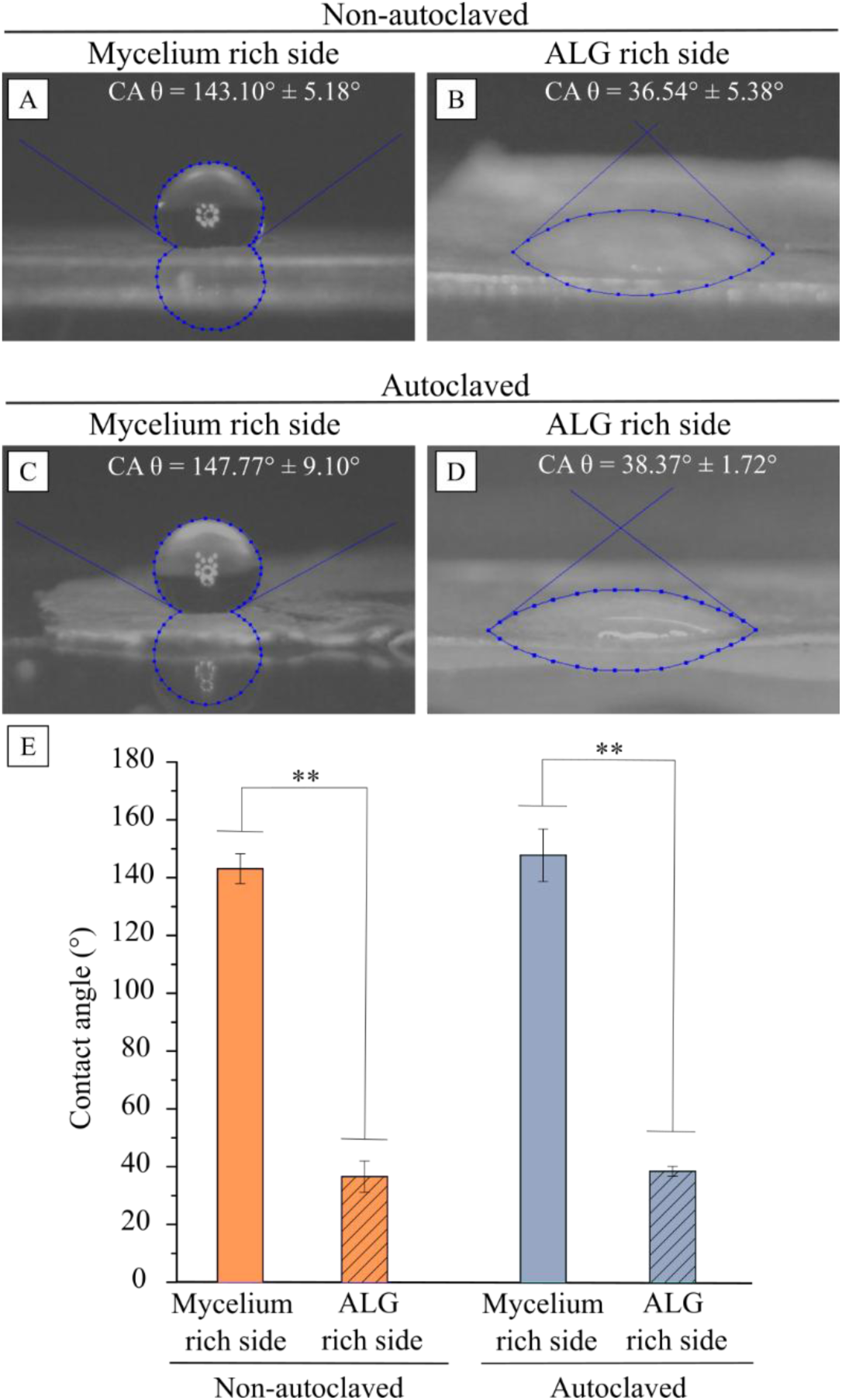
Contact angle measurements tested on biomaterial My-ALG-1.1HA. Photograph of a drop on: A) Mycelium-rich side, and B) ALG-rich side of non-autoclaved biomaterial. C) Mycelium-rich side, and D) ALG-rich side of autoclaved biomaterial. E) Mean contact angle (°) measurements are presented as a bar graph for comparison. The statistical significance is shown as ** p < 0.01. No statistically significant were observed between autoclaved and non-autoclaved groups by Student’s t-test.

Hydrophobic surfaces could be advantageous due to bacterial adhesion inhibition ^42^. Moreover, hydrophobic regions can impart mechanical robustness and stability to the biomaterial structure^43^ while hydrophilic regions facilitate cell-biomaterial interactions and nutrients and waste exchange^44^. Thus, combining hydrophobic and hydrophilic sides within an anisotropic biomaterial could create a more biomimetic microenvironment that better regulates cell behavior and tissue regeneration.

### 3.4 Mechanical and stability properties

The macroscopic analysis of the My-ALG-HA membrane demonstrates an integrated structure that is both facile to handle and flexible (Figure 8A). Additionally, superficial variations between the mycelium-rich side and the ALG-rich side are clearly evident at the macroscopic scale (Figures 8B and C). This observation is consistent with findings from the preceding section concerning surface anisotropy.

**Figure 8.**
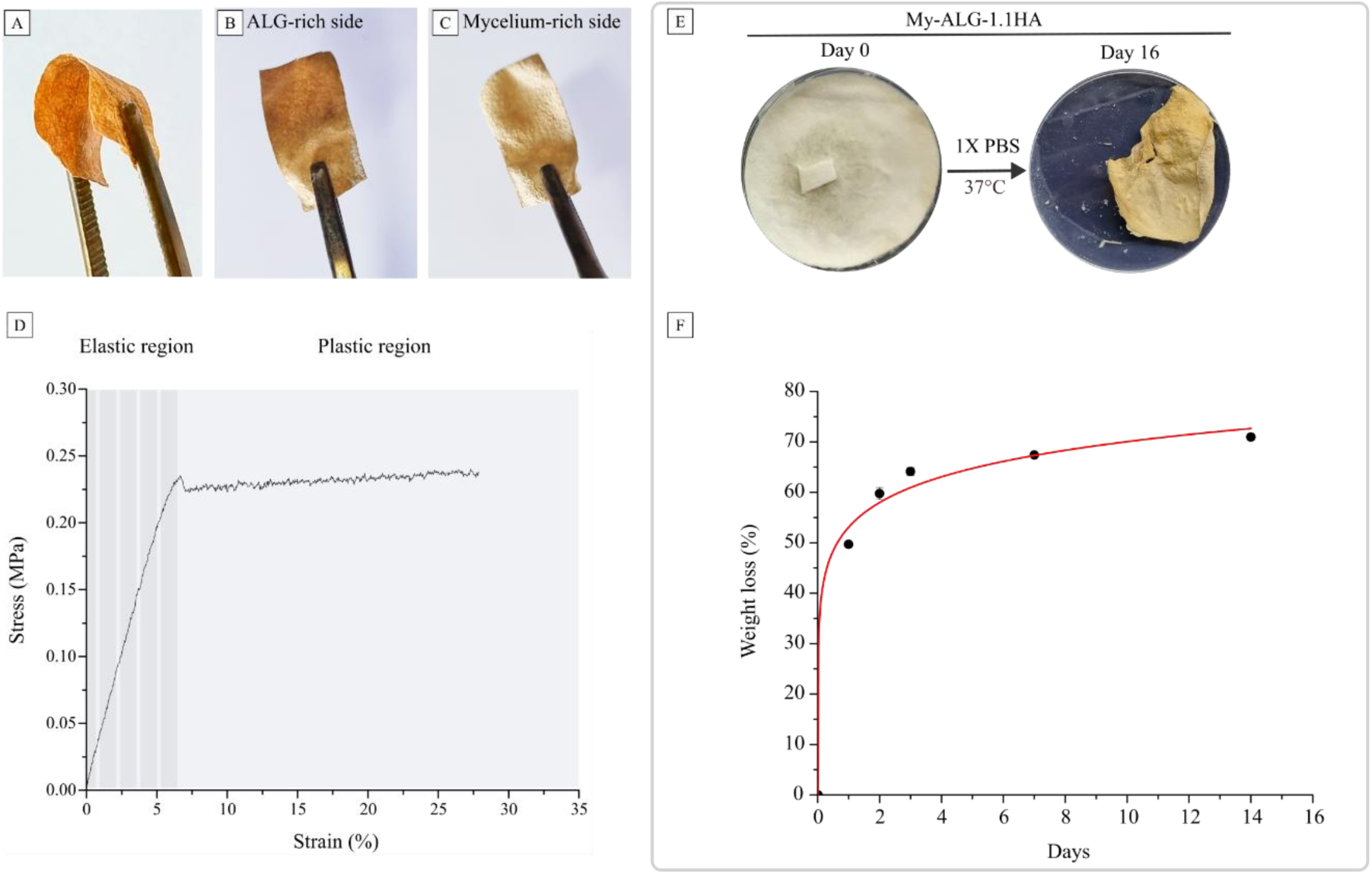
Mechanical and stability properties of the My-ALG-1.1HA membrane. A) Digital photos of the bending membrane, B) macroscopic aspect of the ALG-rich side and C) the mycelium-rich side. D) Representative stress – strain curve. E) Photographic documentation of the biomaterial My-ALG-1.1HA at the beginning and after 16 days of immersion in 1X PBS (pH = 7.4) and 36 °C. F) Weight loss (%) over time.

The mechanical strength of biomaterials must match the requirements of the specific regenerated tissue. For instance, GBR membranes require adequate mechanical support to facilitate bone growth and withstand physiological loads. The design must consider the mechanical properties to ensure the scaffold can support tissue regeneration effectively. Strain-stress curves obtained from the tensile test showed that biomaterial My-ALG-1.1HA has distinct stages: elastic region, yielding and a significant plastic region (Figure 8D).

Values of important parameters such as the elastic modulus and maximum tensile stress, 25.28 ± 1.14 MPa and 0.32 ± 0.07 MPa respectively, were comparable to other mycelium-based biomaterials^14,45^. Interestingly, the maximum tensile strain was 22.20 ± 5.83 % (Table 2), similar to other *Ganoderma* mycelium biomaterial^14^, but higher than mineralized mycelium biomaterials^18,46^. Recent investigations have demonstrated that the tensile strength of oral mucosa tissues varies significantly, ranging from 0.83 MPa to 4.75 MPa, depending on the specific anatomical site. Interestingly, although these values exceed the tensile strength of the My-ALG-1.1HA biomaterial, the Young’s modulus of My-ALG-1.1HA biomaterial is in the range of oral mucosal tissues, 2.22 MPa to 54.72 MPa^47^. This similarity with oral tissues may enhance the biomaterial applicability in GBR, where mimicking the mechanical characteristics of native tissues is crucial for successful integration and functionality.

**Table 2.** Parameters obtained in the tensile assay of the biomaterial My-ALG-1.1HA.

| $\sigma_y$ / Mpa | $\sigma_{max}$ / Mpa | $\epsilon_y$ / % | $\epsilon_{uts}$ / % | $\epsilon_{max}$ / % | E / Mpa |
| --- | --- | --- | --- | --- | --- |
| $0.29 \pm 0.06$ | $0.32 \pm 0.07$ | $6.07 \pm 1.83$ | $19.77 \pm 6.61$ | $22.20 \pm 5.83$ | $25.28 \pm 1.14$ |

Yield stress ( $\sigma_y$ ) and maximum tensile stress ( $\sigma_{max}$ ), yield strain ( $\epsilon_y$ ), tensile strain at the maximum tensile strength ( $\epsilon_{uts}$ ), maximum tensile strain ( $\epsilon_{max}$ ) and Young's modulus (E). Values are expressed as mean ± standard deviation.

In our previous work^18^, we proved the HA contribution to the brittleness of ALG-HA membranes. In this opportunity, the effect of the same amount of HA in M-ALG-1.1HA seems to be masked by the mycelium presence, obtaining a more ductile biomaterial. The constant stress plateau observed in the stress-strain curve of My-ALG-HA after the yield point shows the material’s ability to absorb significant energy before failure. This is a consequence of a material’s crystal structure rearrangement, which increases its toughness. In this sense, ductile biomaterials would be advantageous due to their capability to sustain large strains before fracture, which could help absorb energy and distribute loads in orthopedic applications^48^.

Understanding the degradation rate of biomaterials is essential, as it must be synchronized with the bone healing process to ensure optimal outcomes in bone tissue regeneration. Figure 8F shows the weight loss of My-ALG-1.1HA biomaterial over time in PBS. The weight loss of the biomaterial significantly increased over the first week after immersion. On days 1, 2, and 3, the degradation rates were 49.7 ± 0.6 %, 59.7 ± 1.2 %, and 64.1 ± 0.8 %, respectively. Following day 7 of immersion, the weight started to rise steadily, reaching 67.4 ± 0.6 % by day 7 and 70.9 ± 0.6 % by day 16. Compared to previous findings^18^, the degradation of My-ALG-1.1HA biomaterial was significantly slower than that of ALG-1.1HA membranes, with half of the biomaterial degrading approximately 380 % more slowly. It indicates that incorporating mycelium molecules into this novel biomaterial enhances its hydrophobicity, increasing its resistance to degradation in a simulated physiological environment.

### 3.5 Biocompatibility *in vitro*

To assess hemocompatibility, human blood samples were incubated with 3.4 mg of My-ALG-HA biomaterials. Quantification of hemolysis levels revealed percentages of 1.72 ± 0.26 %, 1.50 ± 0.29 %, 1.77 ± 0.22 %, and 1.93 ± 0.38 % for My-ALG, My-ALG-1.1HA, My-ALG-4.4HA, and My-ALG-8.8HA, respectively (Figure 9). All values were below the 5 % threshold for limited hemolysis, indicating an acceptable hemocompatibility profile. No statistically significant differences were observed with increasing HA concentrations, supporting previous HA hemocompatibility studies^49^ (Figure 9 B).

**Figure 9.**
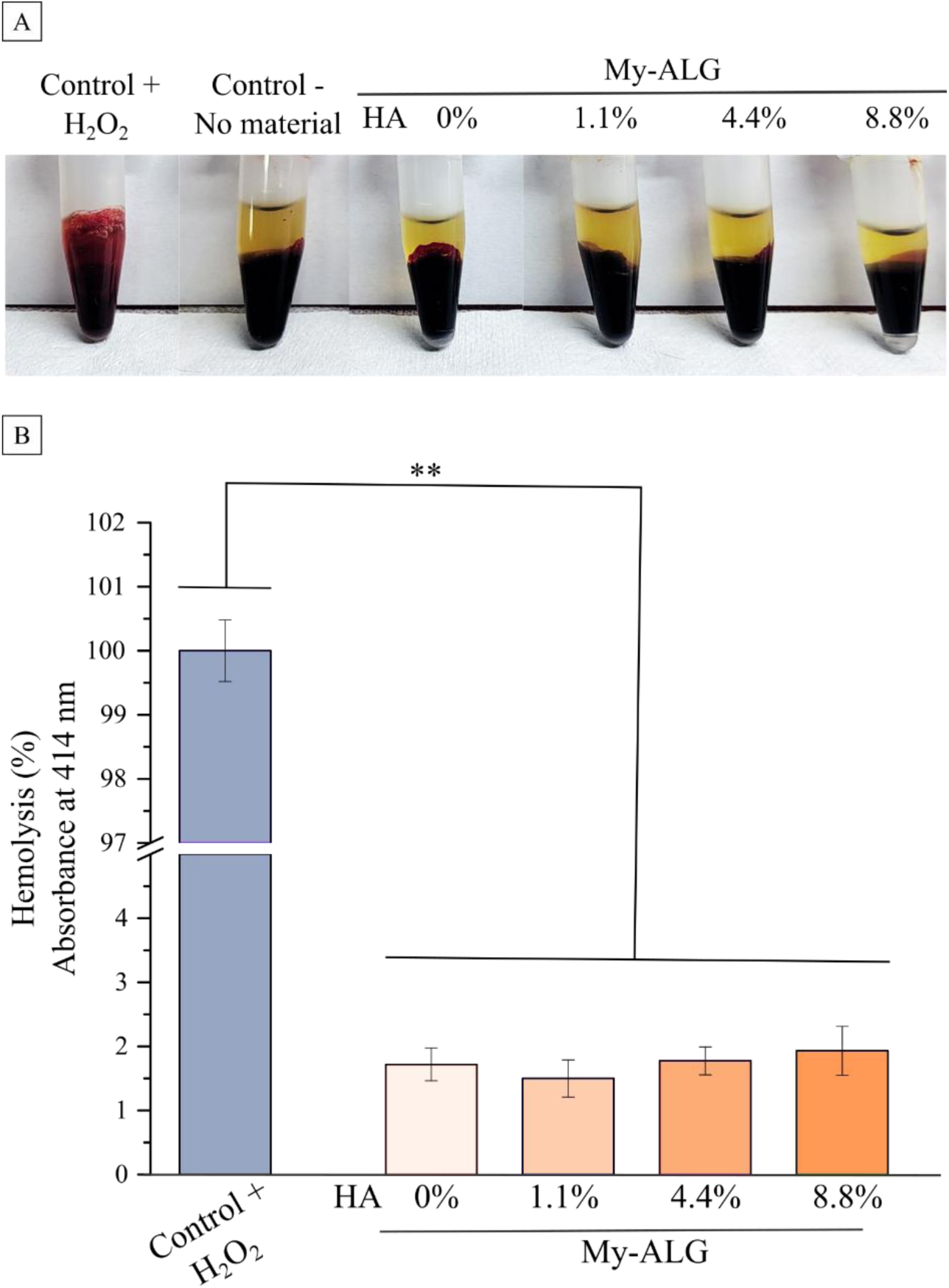
Hemocompatibility assay. A) Photographic documentation of the samples tested immediately before absorbance measurements. B) Hemolysis ratio of human blood in presence of biomaterials My-ALG, My-ALG-1.1HA, My-ALG-4.4HA and My-ALG-8.8HA immersed in 37 °C for 24 h. Hydrogen peroxide solution (H_2_O_2_) was used as positive control. The statistical significance between biomaterials and controls is shown as ** p < 0.01. No statistically significant differences were observed between biomaterials with different HA concentrations by Student’s t-test.

To assess cytocompatibility, MC3T3-E1 osteoblast-like cells were exposed to dispersions of My-ALG, My-ALG-1.1HA, My-ALG-4.4HA, and My-ALG-8.8HA biomaterial powders in complete medium. Cellular morphology was evaluated via optical microscopy after 48 h of incubation. The results indicated that cell size, shape, and spreading were preserved in the presence of the biomaterials, with no observable alterations compared to the untreated control group (Figure 10 A and B). Furthermore, no significant differences in cell viability were detected between cells exposed to different biomaterials and the control group (Figure 10 C). These findings indicate that the biomaterials tested do not negatively affect MC3T3-E1 cell morphology or cytocompatibility under the evaluated conditions.

**Figure 10.**
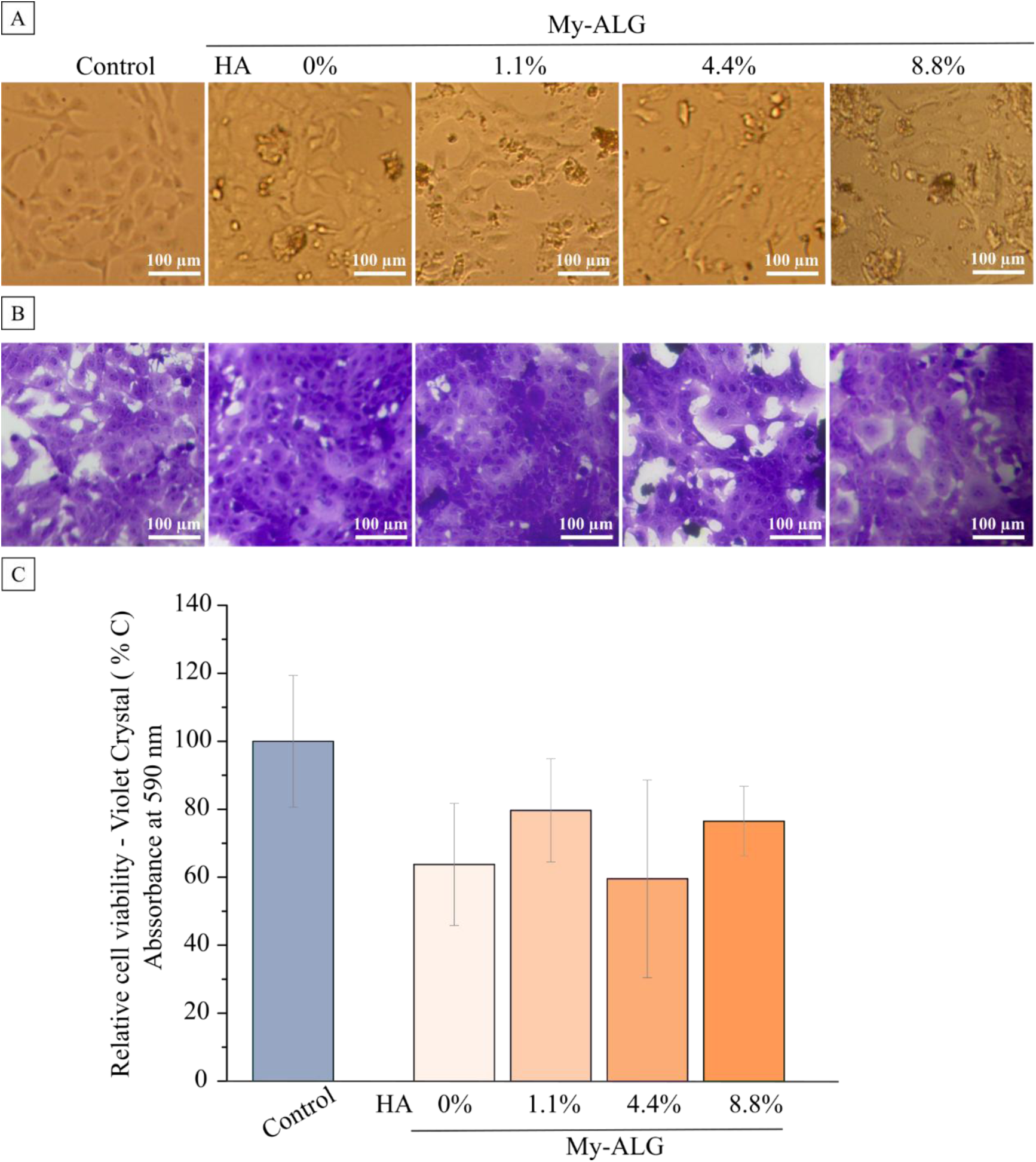
Cytocompatibility assay. A) Inverted light field microscope images showing MC3T3-E1 cells’ morphology and adherence to the surface when cultured in the presence of 1 mg/ml biomaterials My-ALG, My-ALG-1.1HA, My-ALG-4.4HA and My-ALG-8.8HA, before, and B) after crystal violet staining. C) Crystal violet quantification is expressed as relative cell viability (%) in a bar graph. Non-treated cells were used as control. No statistically significant differences were found between the biomaterial conditions and control by Student’s t-test.

## 3 CONCLUSION

This study presented a comprehensive characterization of My-ALG-HA biomaterials, alongside the effective mycelium-inactivation treatments that preserve cell wall integrity and mechanical properties. The mycelium’s successful colonization of the substrate resulted in a trabecular bone-like mineralized network development characterized by a hierarchical fibrillar structure. This structure is particularly notable since the interconnected pores are crucial for nutrients and oxygen supply, as well as the removal of waste. The observed porosity of the biomaterial influences both its mechanical properties and biological performance; thus, it is essential to achieve a balance between them to optimize functionality. Furthermore, the biomaterial’s anisotropic hydrophobic and hydrophilic faces create a biomimetic microenvironment that could enhance proteins adsorption and bone cells attachment while inhibiting bacterial adhesion. Also, mycelium biomolecules in the My-ALG-HA biomaterial enhance its hydrophobicity and resistance to degradation in physiological conditions. Regarding the mechanical properties, its ductile nature allows it to endure significant strains, making it suitable for orthopedic applications where energy absorption and load distribution are critical. The biomaterial maintains favorable osteoblast-like cell morphology, cytocompatibility, and hemocompatibility. While the presence of polysaccharides and triterpenes in *Ganoderma* species suggests therapeutic potential, further research is required to fully characterize the chemical composition and assess the efficacy of these compounds in relevant models. A combination of advanced analytical techniques and targeted biological studies will provide valuable insights into the medicinal properties of *Ganoderma* species. These findings highlight the potential of new mycelium-based biomaterials in advancing bone tissue regeneration, offering innovative insights and opportunities for future developments in the field.

## ASSOCIATED CONTENT

### Supporting Information

Photographic documentation of mycelium inactivation by re-growth assay, X-ray diffraction (XRD) spectrum of the biomaterials, FTIR spectra of the biomaterials, cell wall thickness of the hyphae of the biomaterial obtained by TEM micrographs measurements and histograms of pore sizes distribution of the biomaterial calculated from the SEM images.

## Author Contributions

The manuscript was written through contributions of all authors. All authors have given approval to the final version of the manuscript. ‡PP and JS contributed equally.

## Funding Sources

This work was supported by grants from Agencia Nacional de Promoción Científica y Tecnológica (ANPCyT, PICT-2021-I-A-00108, PICT-2020-3527 and PICT-2020-01922), Consejo Nacional de Investigaciones Científicas y Técnicas (CONICET, PIP-11220210100126CO and PIP-11220210100811CO) and Argentine state on behalf of the ImpaCT.AR program (Impactar D76). D.P. has a postdoctoral fellowship of CONICET, J.S., D.E., V.G.P, N.L.D., P.P. and P.V.M. are researchers of CONICET.

## ACKNOWLEDGMENT

The authors acknowledge to Dr. Virginia Lescano, Dr. Luciano Benedini, Dr. Belén Rauschemberger, Dr. Darío César Gerbino, Techni. Julieta Tornesello Galván and Biol. Cintia Reinoso for their valuable technical contributions.

## ABBREVIATIONS

Backscattered electron scanning microscopy: BSE-SEM
contact angle: CA
Dulbecco′s Modified Eagle′s Medium: DMEM
energy-dispersive X-ray spectroscopy: EDS
fetal bovine serum: FBS
field emission scanning electron microscopy: FE-SEM
Fourier-transform infrared spectroscopy: FTIR
hydroxyapatite nanoparticles: HA
high-Performance Liquid Chromatography: HPLC
malt-yeast extract-sucrose agar: MYSA
maximum tensile stress: σ_max_
mycelium-alginate: My-ALG
mycelium-alginate-nanohydroxyapatite: My-ALG-HA
mycelium-alginate-1.1 % nanohydroxyapatite: My-ALG-1.1HA
mycelium-alginate-4.4 % nanohydroxyapatite: My-ALG-4.4HA) and mycelium-alginate-8.8 % nanohydroxyapatite My-ALG-8.8HA
penicillin/streptomycin: P-S
phosphate buffer: PBS
protein Ling Zhi-8: LZ-8
room temperature: RT
sodium alginate: ALG
Water bidistilled: ddH_2_O
transmission electron microscopy: TEM
tensile strain at the maximum tensile strength: ε_uts_
X-ray diffraction: XRD
yield stress: σ_y_
yield strain: ε_y_
Young’s modulus: E

## Notes

### Competing Interest Statement

The authors have declared no competing interest.

