## Supplementary_Material for "A mycelium biofactory: Novel biomaterial obtained by culturing *Ganoderma sessile* on a potential osteogenic substrate"

[a]Departamento de Química, Universidad Nacional del Sur, Alem 1253, Bahía Blanca 8000, Argentina.

[b]Instituto de Química del Sur, INQUISUR─CONICET, Alem 1253, Bahía Blanca 8000, Argentina.

[c]Laboratorio de Biotecnología de Hongos Comestibles y Medicinales Cerzos, Centro de Recursos Naturales Renovables de la Zona Semiárida (CERZOS-UNS/CONICET), Camino de La Carrindaga Km7, Bahía Blanca 8000, Argentina.

[d]Instituto de Ciencias Biológicas y Biomédicas del Sur (INBIOSUR, UNS-CONICET), Departamento de Biología, Bioquímica y Farmacia, Universidad Nacional del Sur (UNS), San Juan 670, Bahía Blanca 8000, Argentina.

[e] Planta Piloto de Ingeniería Química – PLAPIQUI (UNS-CONICET), Camino La Carrindanga Km 7, Bahía Blanca 8000, Argentina.

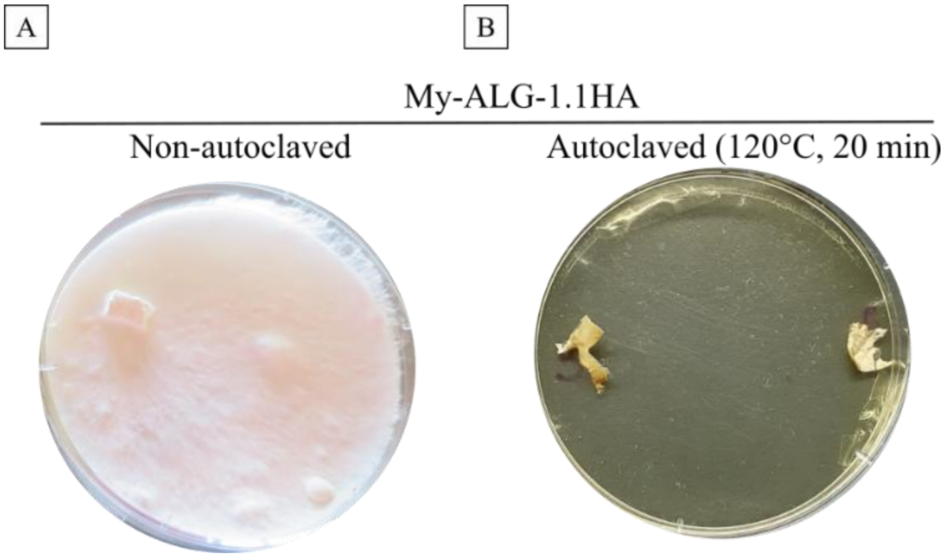


**Figure SM 1.** Mycelium inactivation assay. Photographic documentation of mycelium inactivation by re-growth assay. A) MY-ALG-HA biomaterial without autoclaving, used as control and, B) autoclaved mycelium-based biomaterial MY-ALG-1.1HA. Both portions of mycelia were cultured during 60 days on MYSA medium (2 % w/v malt extract, 0.2 % w/v yeast extract and 1 % w/v sucrose, 2 % agar, pH = 6) at 25 °C.

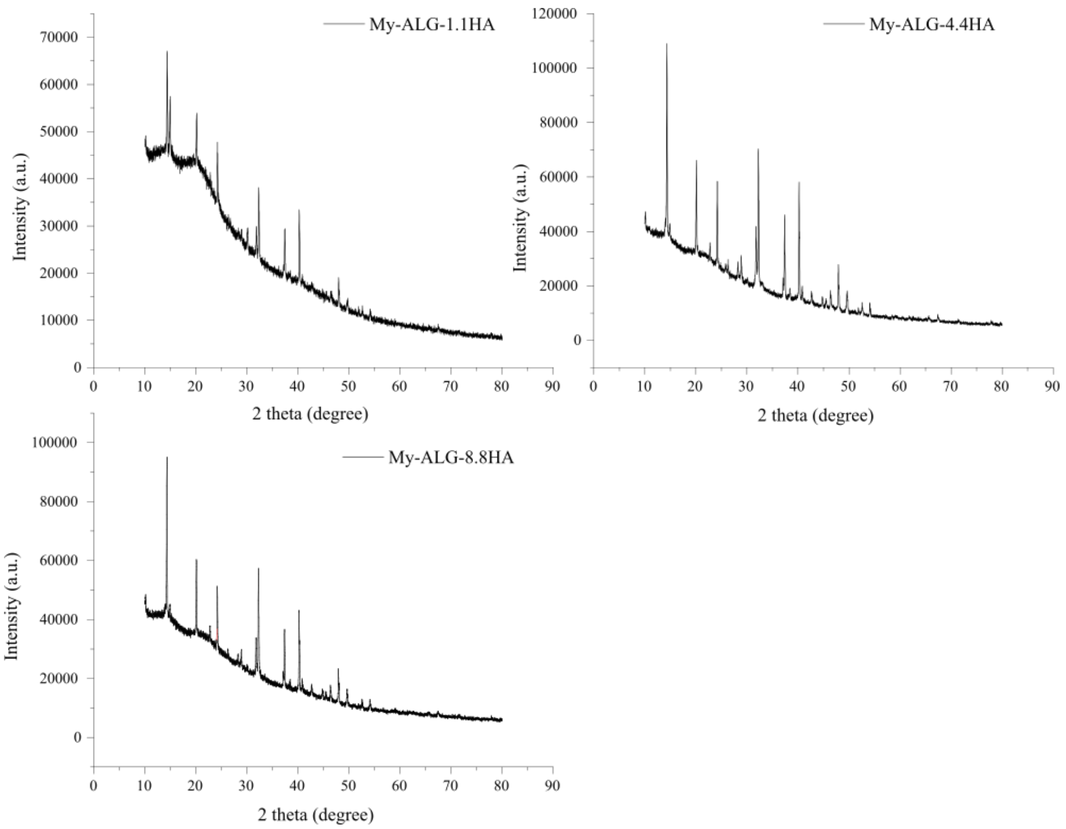


**Figure SM 2.** Chemical composition characterized by X-ray diffraction (XRD) spectra of MY-ALG-1.1HA, MY-ALG-4.4HA and MY-ALG-8.8HA biomaterials. No significant differences were observed among the spectra of biomaterials with varying HA amounts (MY-ALG-1.1HA, MY-ALG-4.4HA and MY-ALG-8.8HA).

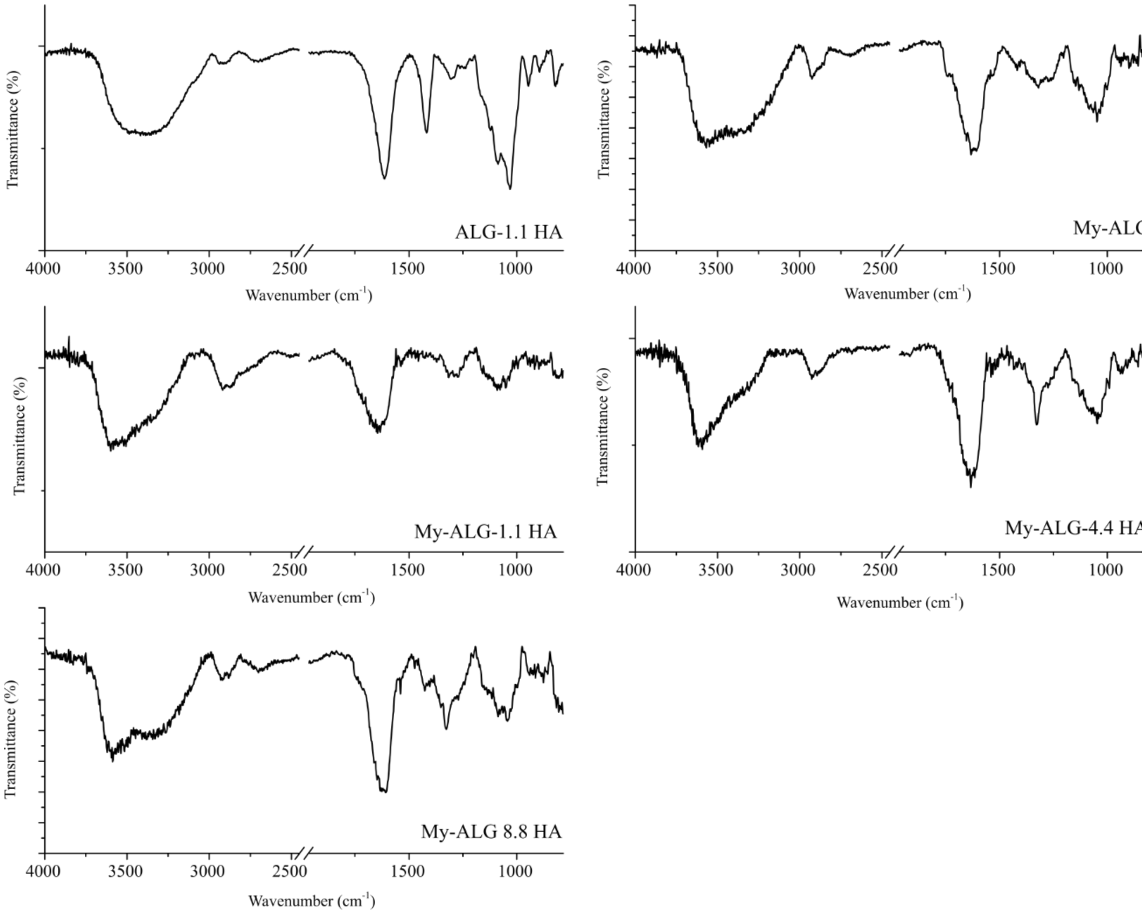


**Figure SM 3**. FTIR spectra of biomaterials ALG-HA, MY-ALG, MY-ALG-1.1HA, MY-ALG-4.4HA and MY-ALG-8.8HA. No differences were found among FTIR spectra of MY-ALG-1.1HA, My-ALG-4.4HA and MY-ALG-8.8HA.

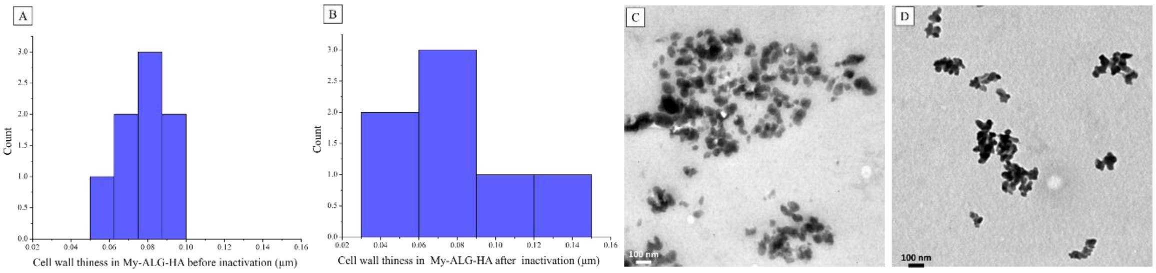

**Figure SM 4.** Cell wall thickness of the hyphae of the biomaterial MY-ALG-1.1HA obtained by TEM micrographs measurements A) before and, B) after inactivation process by autoclaving at 120°C during 20 min. C) Hydroxyapatite nanoparticles present in the My-ALG-HA biomaterial. D) Hydroxyapatite nanoparticles used during synthesis dispersed in water.

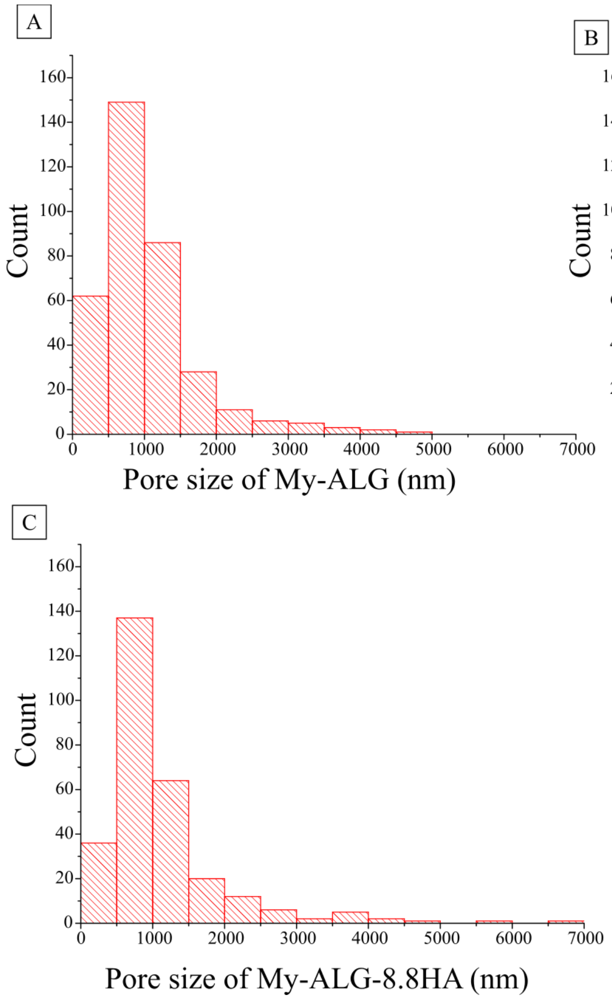


**Figure SM 5.** Histograms of pore sizes distribution calculated from the SEM images of the biomaterials: A) MY-ALG. B) MY-ALG-1.1HA. C) MY-ALG-8.8HA.
